# Complex Dynamics From Simple Cognition: The Primary Ratchet Effect in Animal Culture

**DOI:** 10.1101/134247

**Authors:** Mary Brooke Mcelreath, Christophe Boesch, Hjalmar Kühl, Richard McElreath

## Abstract

It is often observed that human culture, unlike most other animal culture, is *cumulative*: human technology and behavior is more complex than any individual could invent in their own lifetime. Cumulative culture is often explained by appeal to a combination of high-fidelity social learning and innovation, the “ratchet effect.” What is often overlooked is that both human and other animal culture is supported by a more primary ratchet effect that retains and increases the prevalence of adaptive behavior. This primary ratchet can arise without appeal to specialized cognitive adaptations and is plausibly more widespread in animal societies. We use a simple model to highlight how simple forms of contingent social learning can create the primary ratchet effect, dramatically increasing the prevalence of adaptive, hard to invent behavior. We investigate some ways that demography may interact with the primary ratchet to generate patterns of cultural variation. As the primary ratchet may be common to many animal societies, its cognitive components and population dynamics provide a common foundation for the study of animal culture and a necessary foundation for understanding the origins of human cumulative culture.

## 1. Introduction

Culture, defined as socially transmitted behavior, is common in animal societies. Reports of animal culture span a variety of taxa, including mammals, birds, reptiles, fish, and insects (Mundinger 1980, Boyd and Richerson 1996). Next to humans, chimpanzees (*Pan troglodytes*) are thought to have the most prolific repertoire of cultural traditions in the animal kingdom (Whiten et al. 1999, Kühl et al. 2016, Boesch et al. 2016). Orangutans (*Pongo pygmaeus*) also display an impressive array of plausibly cultural traits (Van Schaik et al. 2003). Moreover, captive experiments indicate that many hundreds of species are capable of social learning and culturally transmitted traditions (Laland and Hoppitt 2003).

But human culture remains unusual in at least one respect: it is highly *cumulative*. Much human behavior is more complex than any individual could invent in their lifetime (Boyd and Richerson 1996). It took many generations for relatively simple technologies like bows and baskets to culturally evolve. There is still no consensus about which factors make cumulative behavior possible in humans but largely absent in other animals (Dean et al. 2014). The culture of other animals, including the other great apes, is thought instead to arise from simpler cognitive abilities. There is some evidence that high-fidelity social learning, combined with specialized learning heuristics, can account for cumulative culture (Henrich and McElreath 2003, Henrich and Tennie 2017). Whatever the specific causes, cumulative culture arises from what many authors call the *ratchet effect*: imitation plus innovation allows the population to preserve previous innovations and build complexity across generations (Tomasello 1999, Tennie et al. 2009).

The image that arises from this literature is that human societies are so successful, because of cumulative culture and the specialized individual cognition that makes it possible, while the culture of other animals is largely non-cumulative and of less adaptive consequence (Boyd and Richerson 1995, 1996, Henrich and McElreath 2003). For example, a primary problem with non-cumulative forms of social learning is that they may bring no adaptive benefit at all, despite being easy to evolve (Rogers 1988). Unfortunately, this has obscured the importance of non-cumulative culture.

Nonetheless, social transmission of behavior can be important, even in the absence of cumulative culture. Species without cumulative culture regularly express behavior that is socially transmitted and patterned by population dynamics. In these cases, the distribution of specific behaviors cannot be understood without appeal to cultural evolution. And when the environment changes, predicting how an animal responds will depend crucially upon how it learns. Furthermore, adaptive cultural traditions do not require sophisticated, specialized cognition nor complex, cumulative culture. All they require, in theory, are simple heuristics for the trial and retention of candidate behaviors. These mechanisms may underlie the spread and maintenance of adaptive cultural traditions in many species, including humans.

In this paper, we describe the distinction between the *primary ratchet effect* and the better known *secondary ratchet effect*. We define the primary ratchet as the selective retention of simple, socially transmitted behavior that achieves desired effects and the resort to innovation when no working solution is available. This kind of cultural process is most often described as non-cumulative. There is experimental evidence for strategies of this general type in both human children (Carr et al. 2015) and chimpanzees (Davis et al. 2016). The primary ratchet effect is potentially common to many animals, as it is supported by simple, sequentially structured contingent learning. It allows the spread of solutions to adaptive challenges and can generate diverse patterns of cultural behavior. While the primary ratchet allows simple cultural behaviors to accumulate in terms of their frequency and diversity, the secondary ratchet goes one step further, acting on the traits themselves and allowing cultural behaviors to accumulate enhanced complexity over time.

Even human culture depends upon some of the same cognitive building blocks and population dynamics as the primary ratchet. Many aspects of human culture are simple and possible to invent through individual learning, yet distinct forms are maintained through social learning. Prominent examples include precision throwing and handshaking. Adults use a version of the primary ratchet in laboratory social learning experiments (McElreath et al. 2005), in which social information influences exploration of behavior, but reinforcement learning influences retention and choice. Moreover, the primary ratchet and the cultural accumulation that it generates are relevant to understanding the evolutionary origins of human cumulative culture, which likely arose first from the accumulation of simple traditions before more complex modifications could evolve (Pradhan et al. 2012). In order to understand why non-human animals do not display more cumulative culture, we need a proper origin story for cumulative culture that does not overlook the adaptive benefits of non-cumulative culture.

We present a simple model that demonstrates how contingent learning and the primary ratchet generate non-cumulative culture. This model integrates both individual cognition and population dynamics, demonstrating how the ratchet can produce benefits for both individuals and populations. This general model has been analyzed before, first by Boyd and Richerson (1996) and more extensively by Enquist et al. (2007). We expand the model’s scope to include overlapping generations and subpopulations linked by migration. We explore some of the finite population dynamics and patterns of cultural diversity that such a simple mechanism may generate, showing that some of the demographic properties of cumulative culture, such as a relationship with population density and connectivity (defined here as the extent to which populations are connected through migration) (Shennan 2001, Henrich 2004, Powell et al. 2010, Kline and Boyd 2010, Baldini 2015), are also found in the primary ratchet. However, while demographic and social factors are likely to be important to understanding both ratchet effects, the details could entail important subtle differences.

Our ultimate aim, not yet achieved in this short paper, is to generate theoretical predictions linking models of animal cognition to observational studies of animal culture. This program of research offers a way to link the studies of human and animal culture by exploring the dynamic properties of simple, socially transmitted behavior and highlighting potential homologies across taxonomic groups.

## 2. Contingent learning and the primary ratchet

In this section, we present a model that illustrates how contingent learning can make hard-to-invent behaviors prevalent. In the sections to follow, we incorporate additional demographic factors. We consider a family of learning heuristics, *contingent learning*, whereby individuals sample behavior from other individuals, attempt to achieve a result using the sampled behavior, and then retain the behavior when some desired outcome is achieved. When the sampled behavior is not retained, the individual instead attempts to innovate a new behavior.

### 2.1. Model definition and solution

Assume a large population of organisms capable of only simple, unbiased social learning. Individuals may maintain or reject socially-acquired behavior, based upon subsequent individual experience. Generations are discrete and, for the moment, non-overlapping. Juvenile individuals first learn socially from a random adult, acquiring a candidate behavior from the previous generation. They then try out the behavior and, contingent on cues of its success, either retain the behavior or attempt to innovate a new behavior. Some behavior is adaptive under current conditions, meaning it succeeds at some specific task, such as extracting food, and produces a cue that encourages an individual to retain it. All other behavior is non-adaptive and produces such a cue less often. Conditions change each generation with probability *u*, rendering all previous behavior non-adaptive, which provides an evolutionary incentive to invest in innovation.

A naive individual will encounter an adaptive role model with probability *Q*_*t*_. When this occurs, the individual will either retain the adaptive behavior with probability 1 − *e* or mistakenly reject it with probability *e*. When rejected, individuals will innovate with a probability of success *s*. One can think of the *e* parameter as an error rate in social learning, or the opposite of transmission fidelity *f*=1 − *e* Naive individuals encounter non-adaptive role models with probability 1 − *Q*_*t*_. In this case, there is a chance *d* that an individual correctly rejects the non-adaptive behavior and decides instead to innovate with probability of success, *s*.

With these assumptions, we can write an expression for the frequency, or prevalence, of adaptive behavior in the population at time *t* + 1:

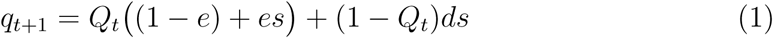

the probability of sampling adaptive behavior, is defined as:

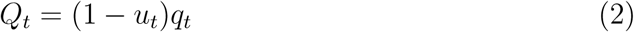

In this expression, *u*_*t*_ = 1 when the environment has just changed between *t* and *t*+1. Otherwise *u*_*t*_ = 0 when the environment has remained the same. This recursion can be solved explicitly for the frequency of successful behavior *T* generations after the most recent change in the environment:

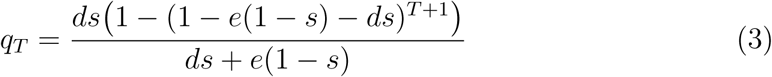

As *T*→∞ reaches a steady state frequency of adaptive behavior 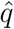 at:

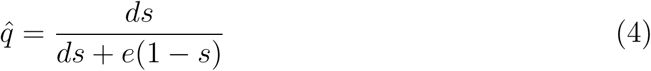

Therefore this is also the maximum prevalence of adaptive behavior that the population can attain. If learning dynamics are much faster than ecological dynamics, then this expression will provide a good approximation of the expected prevalence of adaptive behavior. More generally, since the environment changes in a proportion *u* of the generations, the expected frequency of successful behavior will be lower than 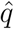. It is found by taking the expectation of (1) with respect to time *t* and solving for the expected value *q*. This yields:

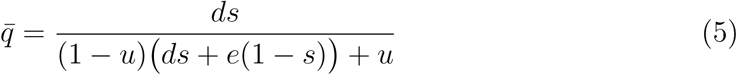

### 2.2. The prevalence of adaptive behavior

With these results, we can address the basic issue of how contingent learning, and the primary ratchet it generates, can dramatically increase the prevalence of adaptive behavior, even when the innovation rate *s* is very small. First, consider the prevalence of adaptive behavior in the absence of social learning. In that case, all naive individuals attempt to innovate, resulting in a frequency of adaptive behavior equal to *s*, the innovation rate. Unbiased, non-contingent social learning will result in the same prevalence, as has been shown in many previous models (Rogers 1988, Boyd and Richerson 1995). The prevalence, *q*, arising from the primary ratchet will exceed *s* as long as:

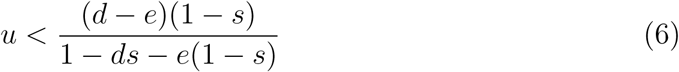

When adaptive and non-adaptive behaviors are always recognized correctly, *d* = 1 and *e* = 0, and then the condition above is always satisfied. For sufficiently large *d* and small *e*, it is trivially satisfied.

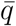 can be many times larger than *s*, provided individuals are good at diagnosing successful techniques. Figure 1, left, shows values of 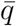 as a function of the innovation rate *s* and rate of environmental change *u*. For small values of *s* especially, the prevalence 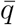 can be several multiples of the innovation rate, even when the environment changes very quickly.

**Figure 1.**
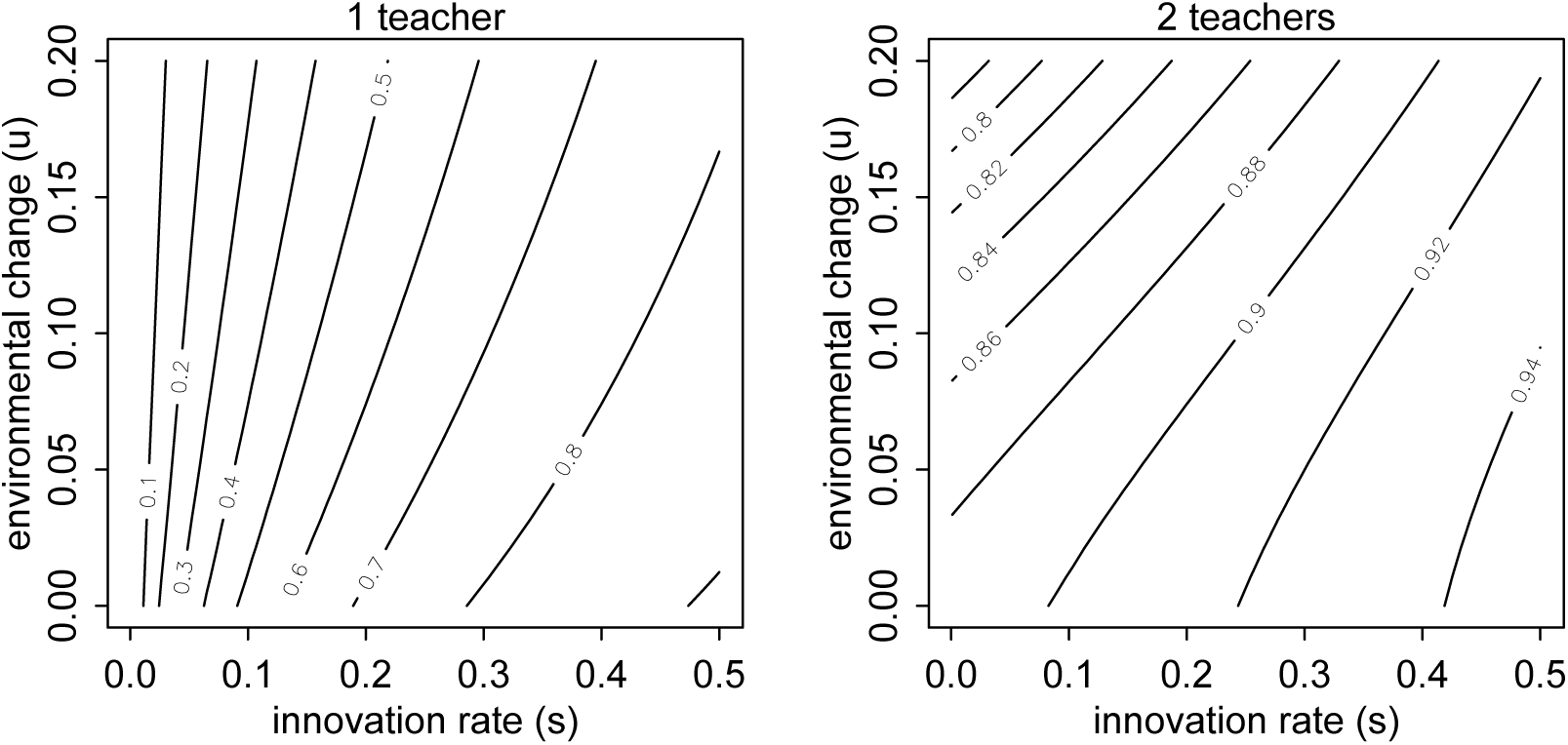
Prevalence of adaptive behavior under the primary ratchet of contingent social learning, as a function of the rate of environmental change *u* and the innovation rate of adaptive behavior *s*. Left: The basic model, with only a single “teacher.” Even at high rates of environment change, the prevalence of adaptive behavior is many times greater than its innovation rate. Right: The general model, with *n* = 2 “teachers.” Prevalence is high, even for very low values of *s*. *d* = 1 and *e*=1/10

The prevalence 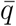 will be even larger, if we allow a simple and plausible modification of *Q*_*t*_, the probability of sampling adaptive behavior. Suppose non-adaptive behavior in a relevant foraging context is just the absence of a solution—individuals who fail to successfully innovate simply do not perform a behavior. This is reasonable for tasks like termite fishing. In this case, it makes little sense that naive individuals would try out the absence of a solution. Suppose instead that each naive individual samples *n* potential “teachers” from a local group. If any one of them displays a successful technique, it can be learned. The only change to the model required here is to redefine *Q*_*t*_:

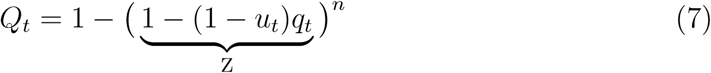

This expression states the probability of sampling at least one teacher with adaptive behavior, out of *n* teachers. The term labeled Z is the probability any of the *n* teachers does not have adaptive behavior. Exponentiating by *n* yields the probably that none have adaptive behavior. Finally, subtracting from 1 yields the probability that any of the teachers has adaptive behavior.

There is no longer a steady state solution for *q*, for general *n*. But solutions can be derived for specific values of *n* or otherwise solved numerically. The right hand plot in Figure 1 shows the values of 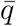 computed using Expression 7 above with *n* = 2. The addition of one more teacher has a dramatic effect on the prevalence of successful behavior. Now even for very low values of *s*, the prevalence of successful behavior is over 0.8.

### 2.3. Evolutionary dynamics

To be a credible candidate for animal social learning, a strategy like contingent social learning should increase the relative fitness of an individual. This is true whether we expect genetic transmission of the strategy or other learning mechanisms to bootstrap the strategy. Our interest is in the behavioral consequences of the primary ratchet. But if it is unlikely to evolve, a reader should be skeptical of its relevance. Boyd and Richerson (1996) have previously shown that a similar strategy readily replaces pure innovation and non-contingent social learning. Here we sketch a proof for our model that reaches the same conclusion.

Suppose an individual who possesses adaptive behavior receives an average fitness increment of *b*. Also suppose that the marginal cost of innovation is *c*. Then assuming weak selection, relative to the cultural time scale, the expected marginal fitness of an individual who only attempts to innovate and never learns socially, is *sb − c*. This is the same expected marginal fitness as a non-contingent social learner, because the fitness of these two simple strategies must be equal at steady state (Rogers 1988). A contingent learner, in contrast, will have marginal fitness:

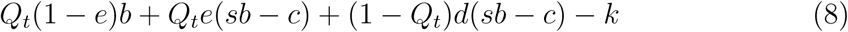

where *k* is the cost of evaluating the efficacy of sampled behavior. This expression is just a sum of all the ways for an individual to acquire adaptive behavior, along with the costs of acquiring it in each case. A contingent learner will have higher relative fitness than a pure innovator or non-contingent social learner, provided:

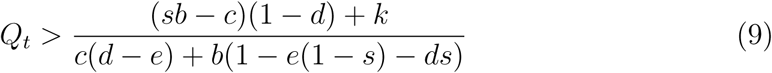

This looks complex, but indicates only that the chance of acquiring adaptive behavior by social learning, *Q*_*t*_, must exceed a ratio of the marginal cost of evaluating behavior to the marginal benefit. As long as *d* is large and the cost of evaluation *k* is small, this condition can be easily satisfied. This is easier to see if we allow *d* = 1 and *e* = 0, simplifying the condition to:

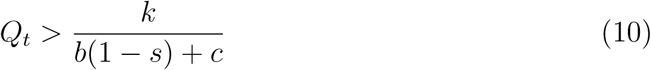

Thus, contingent learning is often superior, provided the cost of evaluation *k* is small, *b* and *c* are large, and *s* is small.

An interesting feature of condition 9 above is that contingent learning, and therefore the primary ratchet effect, can be evolutionarily stable under a wider set of conditions than it can invade a population. This is because the term *Q*_*t*_ will be lower before the primary ratchet effect has increased the prevalence of successful behavior in the population. Once the primary ratchet effect is present, *Q*_*t*_ increases and reinforces the evolutionary advantage of contingent learning. Therefore, as Boyd and Richerson (1996) have emphasized, this type of strategy can be stable under environmental or demographic conditions for which it could not have originally arisen.

### 2.4. Overlapping generations

A conceptual problem with the preceding model is that the parameters *d*, *e*, and *s* integrate the entire lifespan of an individual. Therefore, they are difficult to understand, because presumably more than one learning attempt is possible within a single lifetime—if an individual fails to acquire adaptive behavior in its first year, it can try again in its second. As a consequence, some individuals will require more or fewer attempts to innovate or otherwise socially learn a solution. For example, an individual may initially fail to sample an adaptive behavior, then fail to innovate, and then finally succeed in acquiring adaptive behavior from another individual. In such a case, the model begins to appear internally inconsistent, because the lifetime probability of acquiring adaptive behavior by social learning must be a function of the probability of successful innovation. How can we make sense of the compression of time?

We can begin to unravel this issue by allowing overlapping generations. This means that individuals may live and reproduce for multiple time periods, attempting to acquire or making use of successful behavior in each. Many different contingent learning strategies become possible, with the addition of overlapping generations. For simplicity, we keep the same contingent learning strategy as before, but allow individuals to apply it in each time step. This allows the model to make an important point that is hard to see in the previous model: even quite noisy individual processes in each time step can ratchet up a very high prevalence of adaptive behavior.

Assume that 1 − *μ* is the probability an individual survives from one time period to the next. Population size is regulated such that the number of births each time period equals the number of prevalence of adaptive behavior:

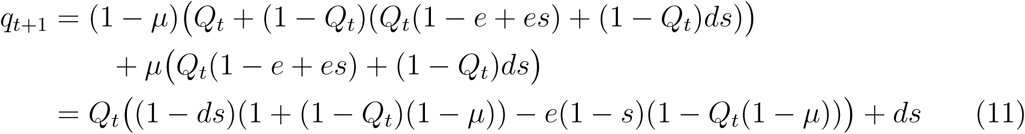

This is very similar to the recursion in the non-overlapping generations model, with the exception that individuals are conservative about adaptive behavior and do not reevaluate it in each time step, until it stops working after a change in the environment. This assumption reflects the notion that *e* is the chance of unsuccessfully applying a technique. Once an individual has successfully learned a technique, the probability *e* does not apply again in each time step. For example, an individual learning for the first time how to fish for termites might have trouble imitating a successful technique and end up rejecting what it has seen. This happens *e* of the time. But once an individual acquires a successful technique, it will only attempt to learn again if the environment changes and renders the technique non-adaptive. As the mortality rateapproaches 1, this model reduces to the previous model with non-overlapping generations.

Expression 11 can be solved for a steady state *q*, but because *q*_*t*+1_ is quadratic in *q*_*t*_, the solution is complicated yielding no direct insight. However, the steady state with overlapping generations will be larger than without, due to the additional accumulation within individual lifetimes. This is most easily seen with a plot similar to those in the previous sections (Figure 2, left). Even computing 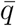 under only a single “teacher” (*n* = 1), the prevalence of adaptive behavior exceeds 0.9 for all combinations of *s* and *u* shown. Higher mortality,μ, reduces the prevalence of adaptive behavior, because asapproaches 1, this model reduces to the model with non-overlapping generations.

**Figure 2.**
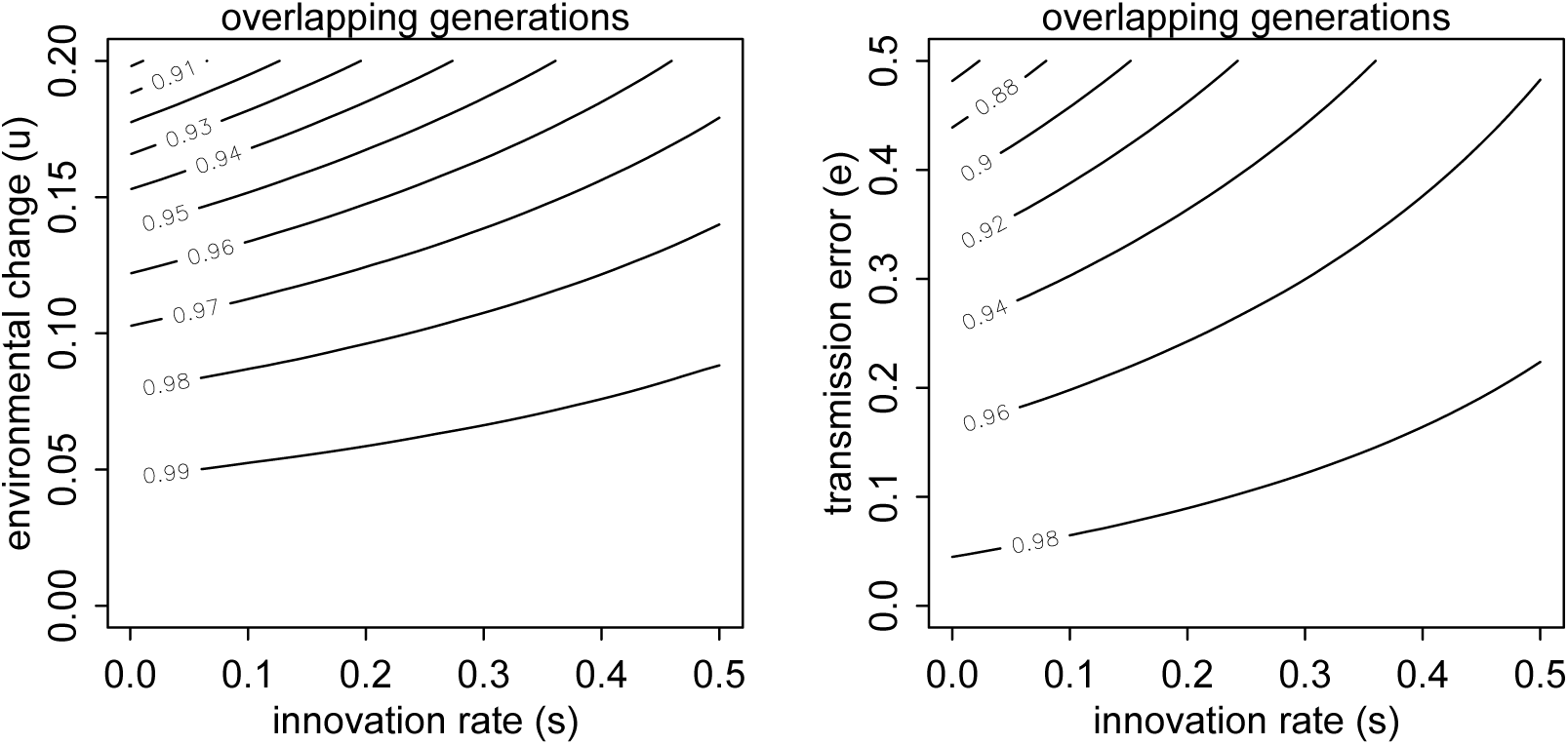
Prevalence of adaptive behavior when generations overlap. Left: Steady state prevalence is now higher than before, for all combinations of *s* and *u*, even here with only a single teacher (*n* = 1). Here shown for *d* = 1, *e* = 1*=*10, and *μ*=0.01. Right: Prevalence is insensitive to the error rate of acquiring adaptive behavior, *e*. Here shown for *u* = 0:1, *d* = 1, and *μ*=0.01

An interesting consequence of overlapping generations and longer individual lifespans is that low-fidelity social learning may not have much impact on the prevalence of adaptive behavior. Figure 2, right, displays 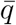 again, but now as a function of the error rate in acquiring successful behavior, *e*, and the innovation rate *s*. The rate of environmental change is set at *u* = 0:1, a rather high rate of change, as environments only stay stable for 10 time steps on average. Nevertheless, prevalence remains above 0.8 for even very high individual error rates.

## 3. Finite population analysis

The results in the previous section address the basic logic and dynamics of the primary ratchet. Contingent strategies that attempt to socially learn available solutions and innovate only when necessary can be individually adaptive and generate a very high prevalence of adaptive behavior. The prevalence values shown in the previous section should not be taken too literally. They are averages, and do not account for the transition periods between environmental changes. They also depend upon a very stylized set of models. But the qualitative results arise from specific mechanisms in the model that depend upon only statistical properties of cognitive strategies. Contingent learning and the primary ratchet can have a substantial impact on animal culture.

In this section we expand the scope of the analysis to individual-based simulations of subdivided populations. The purpose of creating finite groups is to study the influence of group size on the accumulation of socially-transmitted behavior. As mentioned in the introduction, group size and connectivity are thought to influence both behavioral complexity and diversity. We show that the primary ratchet may bear some of the same relationships with demographic structure that are seen in models of cumulative culture. We also investigate patterns of diversity within and between subpopulations, in order to explore patterns that are relevant to studies of animal culture, where many alternative and recognizably distinct solutions to the same problem may be found.

### 3.1. Simulation design

The simulation tracks the behavior of each individual in *G* geographically separated groups, through the sequence of *learning*, *mortality and aging*, *environmental stochasticity*, and *migration*. We focus on a single domain of behavior for which there can be many recognizably distinct adaptive variants. For example, imagine a challenging resource extraction problem such as cracking open a hard fruit or nut—different approaches are possible, and animal cultures sometimes show that local groups vary in which technique is habitually used. The simulation keeps track of “adaptive” variants of behavior—following our example, those that result in successful extraction of the resource—with unique positive integers. All non-adaptive behaviors are coded with zeros, which means that they cannot be distinguished, being absences of solutions.

#### Learning

Learning works in the simulation model exactly as described in the overlapping generations model in the previous section. However, each successful innovation event generates a new, unique behavioral variant that is tracked by a unique identifier. This allows us to track the diffusion of particular innovations and assess diversity among solutions. Some of the diversity may be purely stylistic and nonfunctional. While the functional aspects of a technique may not be very diverse, the addition of non-function steps and postures can make the possible behavior space very large.

#### Mortality and fertility

Groups are regulated by density-dependent mortality, which limits their size around a soft upper limit. Specifically we assume a constant birth rate and an individual death rate *μ* = *μ*_0_ + exp(*KN*_*j*_) − 1 in group *j* with population size *N*_*j*_. K is a parameter that determines how mortality scales with density. This ensures that mortality rises exponentially with local density, stabilizing around an expected population size 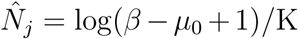. Each time step, the model updates the age of surviving individuals by one year.

#### Environmental stochasticity

Each time step, there is a probability *u* of experiencing environmental change. When environmental change occurs, all solutions in the entire population are rendered non-adaptive.

#### Migration

Groups are also linked by migration, with each individual having a chance *m* of migrating to another random group each time step.

### 3.2. Simulation results

To evaluate the simulation model, we first conducted a broad sweep of parameters, varying *s*, *d*, *e*, *m*,κ, and *u* over wide ranges. We used this opportunity to assure ourselves that we understood the results, on the basis of the underlying analytical model. We provide our simulation code as a supplemental, so that readers can validate and explore the model themselves.

Here, instead of presenting the full sensitivity analysis, we focus on relevant aspects of the simulation that cannot be addressed directly by the analytical model. First, we consider the relative influences of group size *N* and migration rate *m*, a proxy measure of connectivity, as prior work suggests that larger and better connected populations are better able to take advantage of the secondary ratchet effect of cumulative culture. We show here that the primary ratchet effect also benefits from larger groups and high migration rates. Second, we consider how rate of environmental change interacts with the demographic effect of migration. Third, we explore the influence of migration rate *m* as a function of the innovation rate *s*, to demonstrate how much of the interesting behavior of the model avails only at very low values of *s*, where the analogy to cumulative culture is strongest.

In addition to the prevalence of adaptive behavior, we also consider behavioral diversity, as measured by the Shannon diversity index (Shannon 1948). Shannon diversity is the information entropy of the distribution of behavior in the population. We decompose diversity into total diversity and the proportion of diversity between groups, as both are useful ways of quantifying patterns of cultural variation.

In the results, each plotted point is the mean of the final 2000 time steps from each of 10 separate 5000 time step simulations. This duration of simulation was sufficient in all cases to reach steady state. We initialized each simulation at the expected steady state from the analytical version of the model, so convergence to actual steady state was very rapid. In the supplemental, we provide the code needed to reproduce each figure.

#### Group size and connectivity

Both larger and better connected groups achieve higher frequencies of adaptive behavior (Figure 3, left). The smallest groups have the lowest average prevalence of adaptive behavior, for all levels of migration. Migration similarly increases the prevalence of adaptive behavior. The highest migration rate shown, *m* = 0:1, effectively unifies subpopulations and almost entirely cancels any disadvantage of smaller groups.

**Figure 3.**
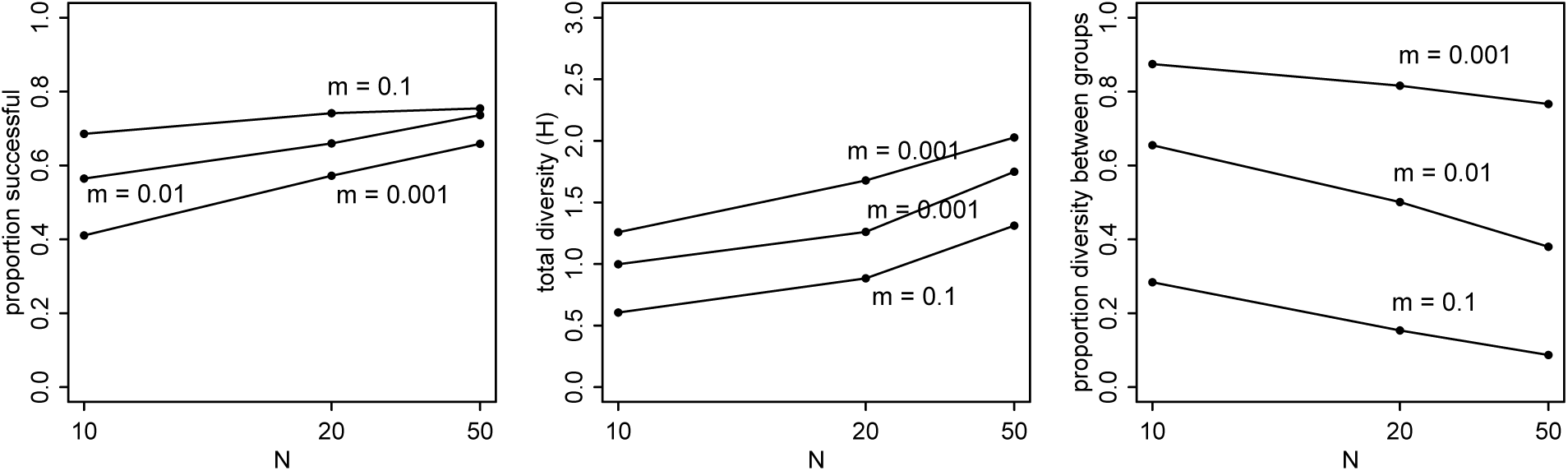
Prevalence of adaptive behavior (left), total behavioral diversity (middle), and proportion of diversity between groups (right) as functions of group size (*N*, horizontal) and migration rate (*m*, separate trend lines). Note inverse scale of horizontal axis, resulting in more linear relationships in plot. Other parameters set to: *s* = 0:001, *u* = 0:01, *μ* = 0:01, *β* = 0:1, *n* = 1.

Note that a migration rate of *m* = 0:1 is very high. The point of these simulations is not to show “realistic” results, but rather to push the system around and understand its forces. The lower migration rate of *m* = 0:001 may be more representative of great ape communities. In this case, smaller local groups suffer quite a lot from the finite population effects.

Note that these results are for a single teacher, *n* = 1. Increasing the number of teachers increases prevalence, as expected. But it does not disrupt the general influence of group size and connectivity.

A consequence of higher prevalence of adaptive behavior is also increased behavioral diversity in the population. Total behavioral diversity (Figure 3, middle) in the population declines both with smaller groups and lower migration rate. However, the effect is reversed when we consider between-group diversity (Figure 3, right). The proportion of diversity between groups is greatest when groups are small and migration is low. These results are as anticipated. It is notable how high the proportion of diversity between groups can be, for reasonable migration rates, such as *m* = 0:001 or even *m* = 0:01 (1% migration per time step). Half or more of the cultural diversity in the population can exist between groups. This result does not require any explicit conformity, just contingent social learning within groups and the action of the primary ratchet effect.

#### Migration and environmental change

When the environment changes rapidly, innovation dynamics are more important, and this makes migration and group connectivity even more important as well. Figure 4 shows again prevalence of adaptive behavior (left), total diversity (middle), and diversity between groups (right) as functions of migration rate (*m*) and the rate of environmental change (*u*). For the very high rate of change, *u* = 0:1 (10 time steps on average between changes in the environment), prevalence and total diversity are suppressed, and migration has only a small effect on either. But for other values of *u*, even small amounts of migration have noticeable impacts of increasing prevalence and decreasing total diversity.

**Figure 4.**
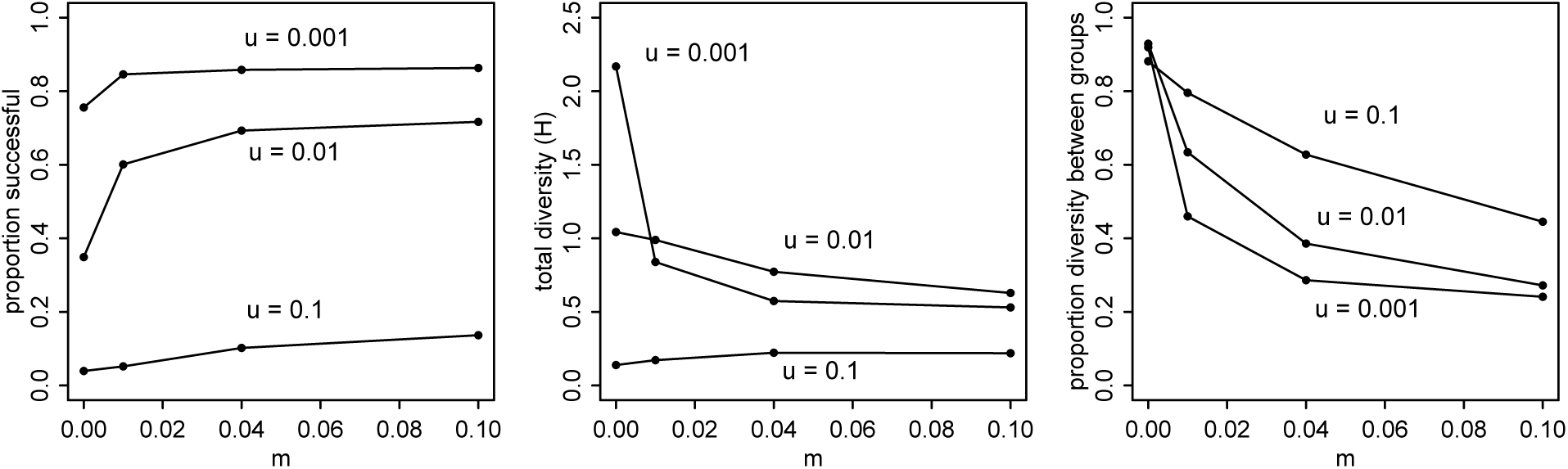
Prevalence of adaptive behavior (left), total behavioral diversity (middle), and proportion of diversity between groups (right) as functions of migration rate (*m*, horizontal) and rate of environmental change (*u*, separate trend lines). Other parameters set to: *s* = 0:001, *N̂* = 10, *μ* = 0:01, *β* = 0:1, *n* = 1.

#### Innovation rate and migration

Finally, we consider variation in innovation rate, *s*. Figure 5 shows simulation results of varying *s* in combination with migration rate, *m*. Note that prevalence of adaptive behavior (left) increases sharply with increases in innovation rate, up to a plateau around *q* = 0:8 where migration rate has only a very small impact. When innovation rates are below about *s* = 0:01, however, population connectivity matters a great deal. This is because when *s* is very small, and groups are well connected, groups can share innovations before the environment changes.

**Figure 5.**
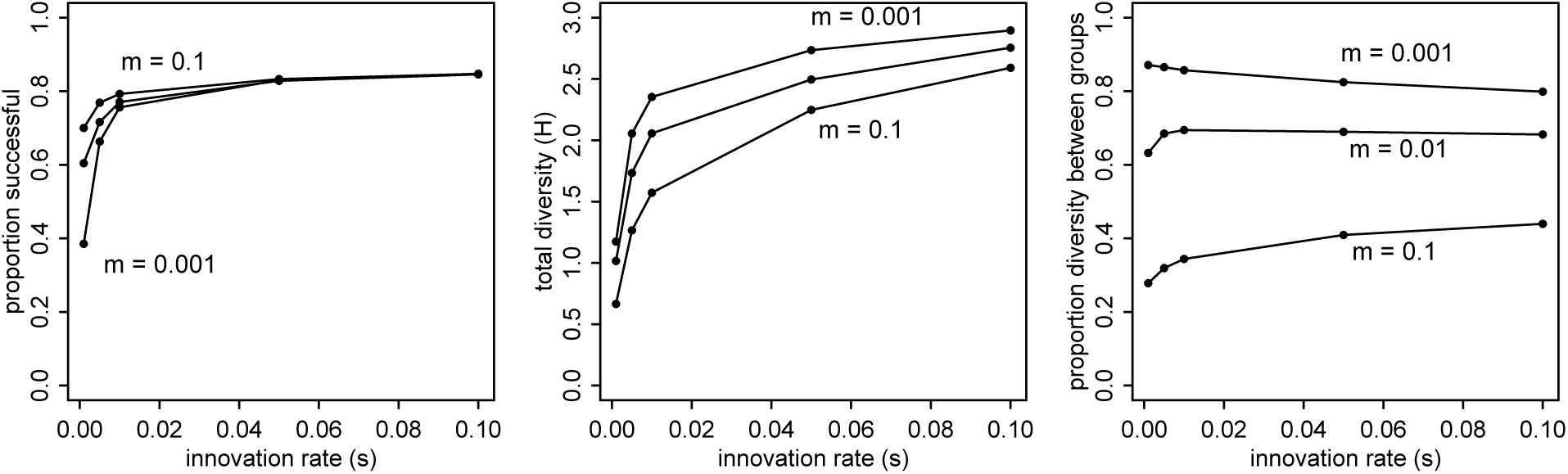
Prevalence of adaptive behavior (left), total behavioral diversity (middle), and proportion of diversity between groups (right) as functions of innovation rate (*s*, horizontal) and migration rate (*m*, separate trend lines). Other parameters set to *s* = 0.001 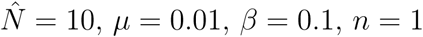

## 4. Discussion

Simple, contingent social learning generates a cultural ratchet effect under a wide variety of demographic conditions. Especially, when innovation rates are very low, the process of contingent learning leads to culturally-transmitted traditions within groups and cultural variation among groups. This result is important, because when *s* is small, successful techniques are difficult for individuals to invent within their lifespans. However, the primary ratchet effect can quickly spread and retain rare innovations. This produces relationships with demography that are quite similar to those found in models of cumulative culture.

In many cases, the effect of the primary ratchet can be distinguished from individual learning, because it generates frequencies of adaptive behavior that are much higher than their underlying innovation rates. Tennie et al. (2009) argue for example that much ape behavior exists in a “zone of latent solutions” that individuals can reinvent, given materials and motivation. Overall prevalence and diversity between groups may still owe a lot to cultural transmission. Thus, it may be possible to identify cultural transmission in the wild based on prevalence data, without requiring direct observational evidence which can be extremely difficult to obtain. This depends upon having some sense of the underlying innovation rate. Limited available evidence suggests that innovation rates are quite low in nature (Perry et al. 2017, Reader et al. 2016). Another option is to exploit age structure and attend to which age classes innovate as well as the prevalence of behavior in all age classes.

The difficulty in obtaining direct evidence of socially transmitted culture in the wild has led some to rely on the “exclusion approach,” whereby cultural processes are inferred by excluding ecological and genetic explanations. This approach emphasizes cultural differences, or high diversity, between groups when environmental, ecological, and genetic differences are minimal. However, our modeling results show that the primary ratchet can generate both high and low levels of behavioral diversity between groups. Thus, cultural similarities between groups should not be interpreted as a lack of evidence for culture, as implied by the exclusion approach. Our results show that, regardless of how behavioral diversity is partitioned among groups, when the prevalence of adaptive behavior far exceeds its innovation rate, cultural transmission is involved.

We additionally found that high-fidelity social learning (*e*), was not required for the primary ratchet to function. In Figure 2 we showed how overlapping generations and extended lifespans make the error rate of individual transmission events poorly representative of the population process. Even when transmission error was as high as 50%, the primary ratchet could raise the prevalence of adaptive behavior over 80%.

This casts doubt on our ability to extrapolate about animal culture from short-term, individual captive experiments, such as those cited by Henrich and Tennie (2017), in which animals display low-fidelity social learning. Notably, there is substantial evidence that social learning in humans can be highly error prone, as well, and many anthropologists suspect that the stability of human cultural traditions has less to do with the accuracy of imitation than is traditionally believed (Sperber 1996).

While this model considers only a single domain of behavior, such as solutions to foraging a particular resource, it is relevant to understanding the accumulation of cultural solutions in a number of domains. Both the empirical analysis of cumulative culture by Kline and Boyd (2010) and the model of cumulative culture by Baldini (2015) actually assume discrete, non-cumulative items and consider how the accumulation of tools or solutions in a number of domains is related to demography. With respect to our model, as long as all domains are independent, then the expected accumulation will be the number of domains *D* times the expected prevalence *q̄*. This is very similar to the results of Baldini (2015), but here in the absence of payoff-biased social learning. It is likely, however, that different domains are not independent, with both positive and negative externalities flowing among them.

The age and mortality structure we assume is very simple—mortality is constant across ages and unconditional on behavior. This may generate odd artifacts, like a long tail of ancient individuals. In this circumstance, an adaptive heuristic could use teacher age to great benefit.

There are many processes omitted from our model that could generate additional cultural dynamics. Adaptive social learning biases like conformity and payoff-bias (Boyd and Richerson 1985, Henrich and McElreath 2003) can further ratchet up the prevalence of adaptive behavior, even in the absence of cumulative complexity. Our point is not to exclude these processes from consideration. Rather, our point is that appeal to such strategies is not necessary for cultural evolution to produce group benefits and be empirically distinguishable from individual learning or the simplest, non-adaptive forms of social learning.

## Supplemental materials

See the repository at: https://github.com/rmcelreath/contingent_learning_finite_pop_sims.

## Acknowledgments

We thank Bret Beheim, Paolo Gratton, and the PanAf Research Group for helpful discussions and comments on this work.

